# Antioxidants green tea extract and nordihydroguaiaretic acid confer species and strain specific lifespan and health effects in *Caenorhabditis* nematodes

**DOI:** 10.1101/2021.11.09.464847

**Authors:** Stephen A. Banse, Christine A. Sedore, Erik Johnson, Anna L. Coleman-Hulbert, Brian Onken, David Hall, E. Grace Jackson, Phu Huynh, Anna C. Foulger, Suzhen Guo, Theo Garrett, Jian Xue, Delaney Inman, Mackenzie L. Morshead, W. Todd Plummer, Esteban Chen, Dipa Bhaumik, Michelle K. Chen, Girish Harinath, Manish Chamoli, Rose P. Quinn, Ron Falkowski, Daniel Edgar, Madeline O. Schmidt, Mark Lucanic, Max Guo, Monica Driscoll, Gordon J. Lithgow, Patrick C. Phillips

**Affiliations:** Institute of Ecology and Evolution, University of Oregon, Eugene, OR 97403, USA; Rutgers University, Dept. of Molecular Biology and Biochemistry, Piscataway, NJ 08854, USA; The Buck Institute for Research on Aging, Novato, CA 94945, USA; Division of Aging Biology, The National Institute on Aging, Bethesda, MD 20892, USA

**Author notes:** Corresponding authors’ e-mail addresses.

## Abstract

The *Caenorhabditis* Intervention Testing Program (CITP) is an NIH-funded research consortium of investigators who conduct analyses at three independent sites to identify chemical interventions that reproducibly promote health and lifespan in a robust manner. The founding principle of the CITP is that compounds with positive effects across a genetically diverse panel of *Caenorhabditis* species and strains are likely engaging conserved biochemical pathways to exert their effects. As such, interventions that are broadly efficacious might be considered prominent compounds for translation for pre-clinical research and human clinical applications. Here, we report results generated using a recently streamlined pipeline approach for the evaluation of the effects of chemical compounds on lifespan and health. We studied five compounds previously shown to extend *C. elegans* lifespan or thought to promote mammalian health: 17α-estradiol, acarbose, green tea extract, nordihydroguaiaretic acid, and rapamycin. We found that green tea extract and nordihydroguaiaretic acid extend *Caenorhabditis* lifespan in a species-specific manner. Additionally, these two antioxidants conferred assay-specific effects in some studies—for example, decreasing survival for certain genetic backgrounds in manual survival assays in contrast with extended lifespan as assayed using automated *C. elegans* Lifespan Machines. We also observed that GTE and NDGA impact on older adult mobility capacity is dependent on genetic background, and that GTE reduces oxidative stress resistance in some *Caenorhabditis* strains. Overall, our analysis of the five compounds supports the general idea that genetic background and assay type can influence lifespan and health effects of compounds, and underscores that lifespan and health can be uncoupled by chemical interventions.

## Introduction

Elaboration of clinical strategies that delay health declines associated with aging is anticipated to markedly improve life quality for the elderly and their families. One research area focused on such a goal seeks to identify compound interventions that can prolong life and/or promote healthy aging. Toward this end, model organisms are invaluable in initial compound screening as these models offer low expense, ease of culture, short lifespans and often simple health assays. One of the most widely studied aging models, *Caenorhabditis elegans,* has proven a useful system for identifying and characterizing compounds that robustly extend lifespan and healthspan^1–3^.

The *Caenorhabditis* Intervention Testing Program (CITP), an NIH-funded research consortium consisting of investigators at three independent sites (Rutgers University, the University of Oregon, and the Buck Institute for Research on Aging), is tasked to identify pharmacological interventions with the potential to extend *Caenorhabditis* lifespan and healthspan in a robust manner. The founding principle for the CITP is that compounds with positive effects across a genetically diverse population engage conserved biochemical pathways that promote healthy aging. The CITP is distinctive in testing compounds across a genetically diverse panel of *Caenorhabditis* strains and species to identify interventions that promote lifespan extension independent of genetic background. Another distinctive feature of the CITP effort is that studies are replicated as closely as possible at the three geographically distinct sites. To date, the CITP has reported on the lifespan and healthspan effects of more than 27 compounds in more than 250,000 individuals across nearly 280 trials^4–9^.

The initial CITP studies focused on longevity as the sole endpoint for anti-aging intervention evaluation^4–7^. For the current suite of test compounds, we expanded and modified our workflow^4^ to evaluate the potential of compounds to extend lifespan and/or healthspan across *Caenorhabditis* species and strains (Supplemental Figure 1).

Here, we report results from five compounds in the CITP testing pipeline: 17α-estradiol, acarbose, green tea extract (GTE), nordihydroguaiaretic acid (NDGA), and rapamycin. 17α-estradiol, a weak endogenous steroidal estrogen, has been reported to alleviate age-related metabolic dysfunction and inflammation in male mice^10^, to protect against neurodegeneration in cell and animal models of Parkinson’s disease^11^, and to extend lifespan in genetically heterogenous male mice^12,13^. Acarbose, an anti-diabetic drug that inhibits alpha-glucosidase, has been shown to prevent age-related glucose intolerance^14^ and to limit postprandial hyperglycemia in mice^15,16^. Likewise, acarbose has been shown to extend median lifespan in genetically heterogenous mice^12,13^. GTE is rich in antioxidant polyphenols, and has been reported to reduce the risk of coronary heart disease and certain forms of cancer^17^, as well as to provide neuroprotection against diseases such as Alzheimer’s^18^. GTE increases lifespan in both flies^19,20^ and mice^21^, and its primary constituent flavonoid, epigallocatechin gallate (EGCG), has been shown to extend mean lifespan in *C. elegans^22^,* although that result may be context dependent^23,24^. NDGA, a lignin found in the creosote bush, possesses both antioxidant and anti-inflammatory properties^25,26^, and has been shown to increase lifespan in mice^12,27^. Finally, rapamycin, an mTOR kinase inhibitor, has been shown to increase lifespan in a variety of model organisms, including mice and flies^28–30^.

Our studies of 17α-estradiol, acarbose, GTE, NDGA, and rapamycin underscore the complexities of assessing biological outcomes of candidate lifespan extending treatments. Using our standardized protocols, we find that the antioxidants GTE and NDGA extend *Caenorhabditis* lifespan in a species-specific manner. GTE and NDGA tests also revealed some assay-specific outcomes—in certain genetic backgrounds we found decreased survival in manual longevity assays, whereas we measured extended lifespan when we determined outcomes using the automated *C. elegans* Lifespan Machines (ALM). GTE and NDGA affected swimming ability in a strain-specific manner, and GTE lowered oxidative stress resistance in some *Caenorhabditis* strains. Lifespan and healthspan could thus be uncoupled as evaluated by our approach, with outcomes influenced by genetic background. Overall, our findings on this test set of interventions underscore how impactful genetic background, selected health assay, and protocol details are in the assessment of intervention effects. Interventions that meet the high bar of efficacy across a broad range of genetic backgrounds and across multiple experimental approaches may prove the exception but would establish definitive priority for testing in mammalian models.

## Results

### The CITP pipeline for the evaluation of compounds on lifespan and healthspan

We created a workflow plan to evaluate the ability of a chemical intervention to extend lifespan and/or healthspan across *Caenorhabditis* species and strains (Supplemental Figure 1). The CITP pipeline involves preliminary testing in a single lab to identify a potentially effective dosage range (phase one study; Supplemental Figure 2) and to target a bioactive dose across strains (phase two; Supplemental Figure 3) before we undertake survival assays in all three CITP labs (phase three; Figure 1). In phase four, we evaluate the effect of bioactive compounds that passed the preliminary tests for robust impact on longevity (phases 1-3) on select health measures (Figures 2 and 3). We publish our findings on compounds without positive lifespan that exit the pipeline^6–8^ (Supplemental Figure 1).

**Figure 1:**
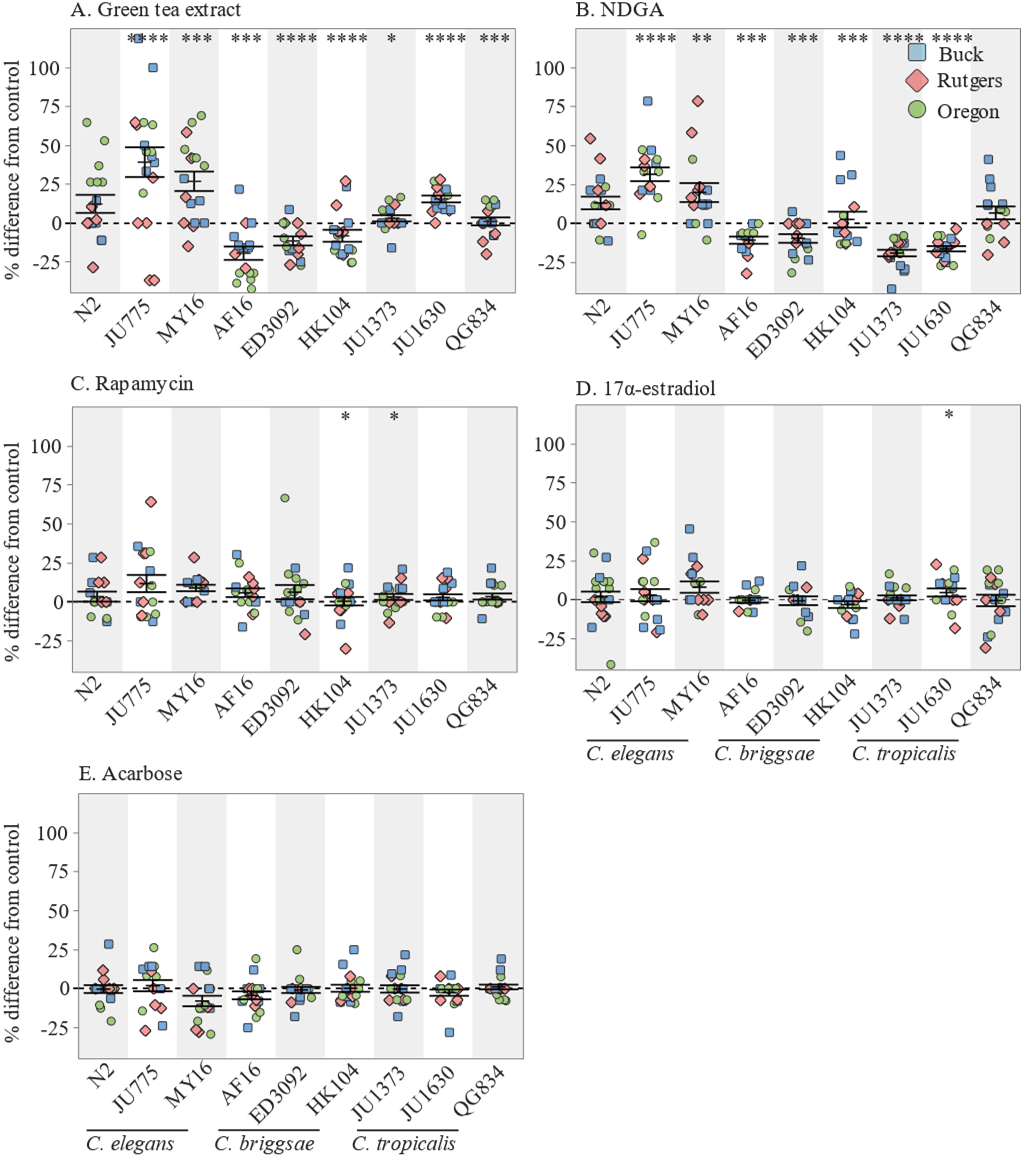
The effect of adult exposure to (a) 500 μg/mL green tea extract, (b) 100 μM NDGA, (c) 50 μM rapamycin, (d) 30 μM 17α-estradiol, and (e) 500 μM acarbose on the median lifespan of nine strains across three *Caenorhabditis* species. Three strains were tested from each species: *C. elegans* N2, MY16, and JU775 (black text), *C. briggsae* AF16, ED3092, and HK104 (dark gray text), *C. tropicalis* JU1373, JU1630, and QG834 (light gray text). Each point represents the change in median lifespan from an individual trial plate (compound treated) relative to the specific control (vehicle only) conducted. The bars represent the mean +/− the standard error of the mean. Replicates were completed at the three CITP testing sites (blue square–Buck Institute, green circle–Oregon, and salmon diamond–Rutgers). Asterisks represent *p*-values from the CPH model such that \*\*\*\**p*<0.0001, \*\*\**p*<0.001, \*\**p*<0.01, and \**p*<0.05.

**Figure 2:**
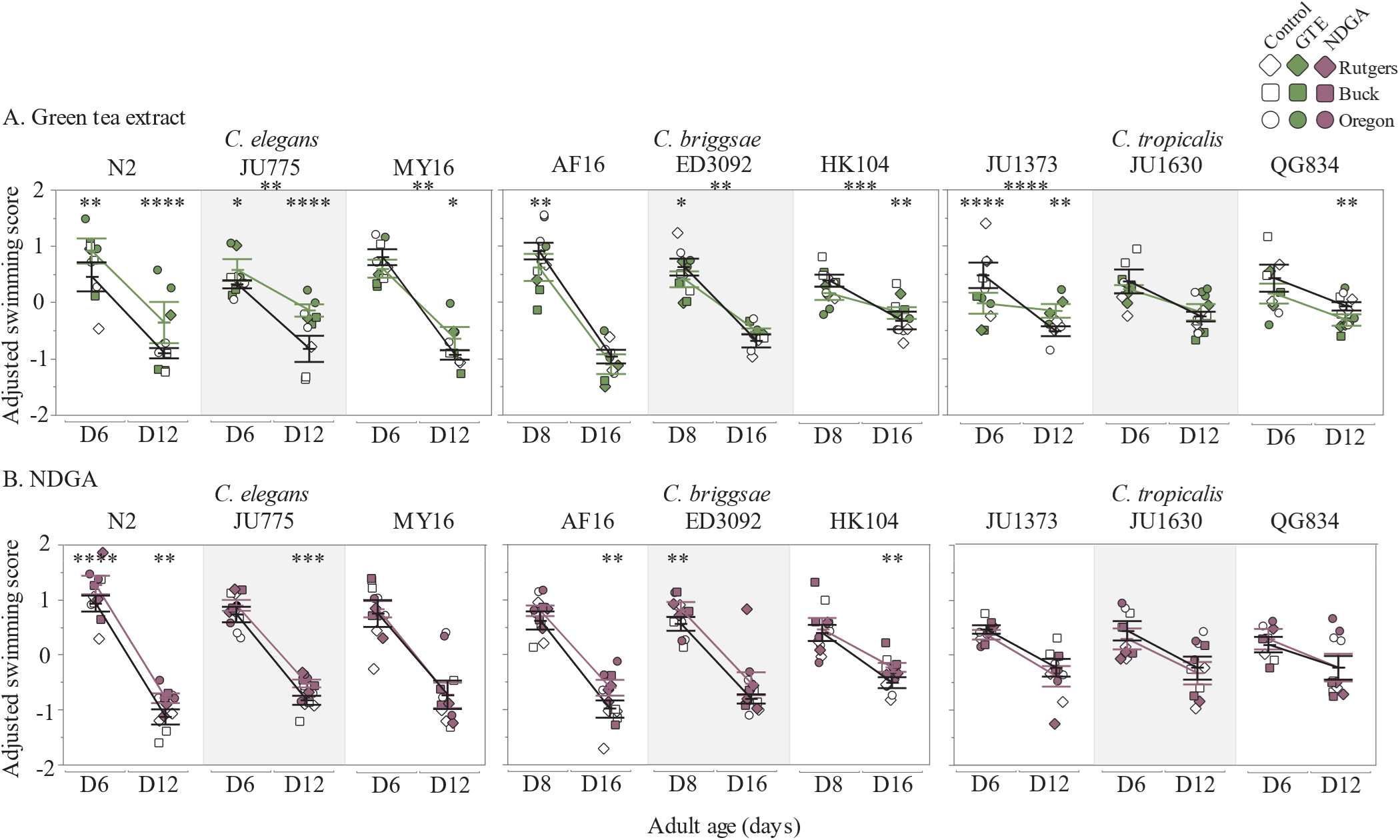
The mean adjusted swimming score across *Caenorhabditis* species after treatment with (a) GTE, or (b) NDGA is shown for young and old ages (adult days 6 and 12 for *C. elegans* and *C. tropicalis,* adult days 8 and 16 for *C. briggsae*). Strains tested include: three *C. elegans* strains (N2, JU775, MY16), three *C. briggsae* strains (AF16, ED3092, HK104), and three *C. tropicalis* strains (JU1373, JU1630, QG834). Each data point represents the mean from one trial, the bars represent the mean +/− the standard error of the mean, and the colors correspond to the treatment conditions starting at day 1 of adulthood (white-DMSO control, green–500 ug/mL GTE, mauve–100 uM NDGA). Replicates were completed at the three CITP testing sites (square-Buck Institute, diamond-Rutgers, circle-Oregon). Asterisks inside the plots represent *p*-values from the linear mixed model, and asterisks outside the plot represent *p*-values from the type III ANOVA indicating a significant compound by age interaction, such that \*\*\*\**p*<0.0001, \*\*\**p*<0.001, \*\**p*<0.01, and \**p*<0.05.

**Figure 3:**
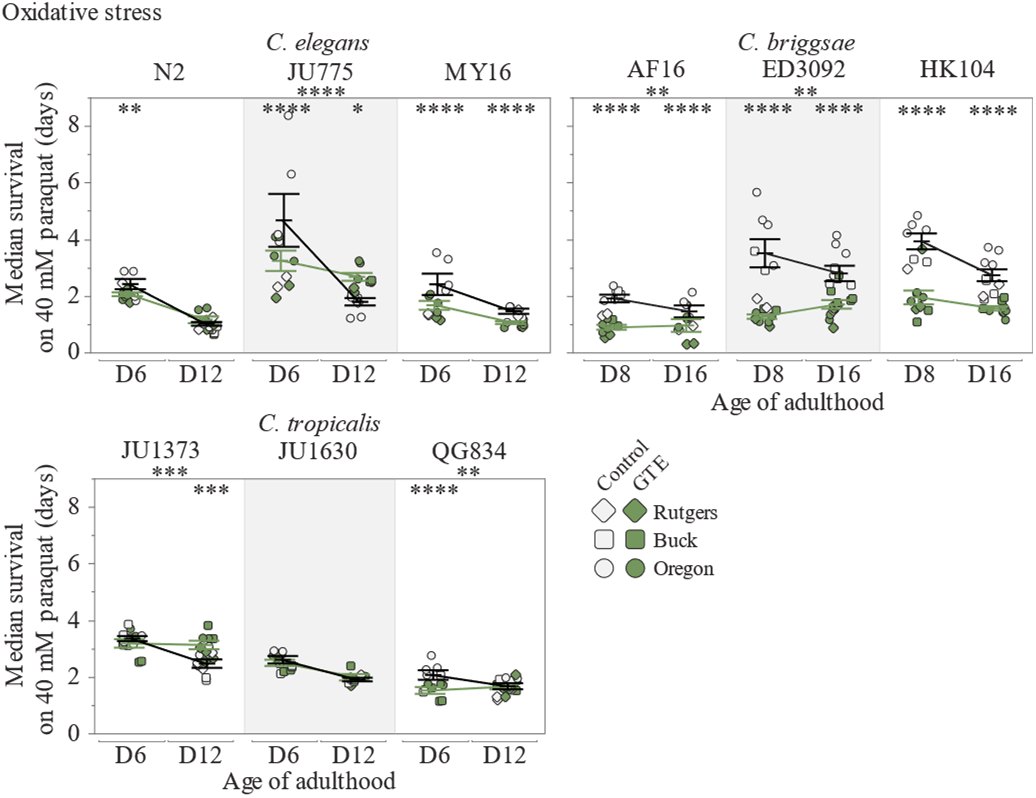
Survival of nine *Caenorhabditis* strains on 40 mM paraquat after adult exposure to green tea extract (green symbols), or vehicle control (open symbols). Oxidative stress resistance was measured at day 6 and 12 of adulthood (*C. elegans* and *C. tropicalis*), or day 8 and 16 of adulthood (*C. briggsae*). Three strains were tested from each species: *C. elegans* N2, JU775 and MY16, *C. briggsae* AF16, ED3092 and HK104, and *C. tropicalis* JU1373, JU1630 and QG834. Dots represent the median survival of one plate replicate (gray–vehicle only control, green–500 μg/mL GTE). Replicates were completed at three CITP testing sites (circle–Oregon, square–Buck, diamond– Rutgers). Asterisks inside the plots represent *p*-values from the CPH model, and asterisks outside the plot represent *p*-values from the type III ANOVA indicating a significant compound by age interaction, such that \*\*\*\**p*<0.0001, \*\*\**p*<0.001, \*\**p*<0.01, and \**p*<0.05.

### The antioxidants green tea extract and NDGA extend Caenorhabditis lifespan in a species-specific manner

Using the streamlined pipeline we describe above, we evaluated a test set of compounds for lifespan effects. Among this test set, we found that rapamycin, acarbose, and 17α-estradiol were broadly ineffectual. Rapamycin only altered lifespan for two (*C. briggsae* HK104 and JU1373) of the nine tested strains, while 17α-estradiol only had an effect on JU1630 (Figure 1). We also found that acarbose did not statistically change the lifespan of any of the tested strains.

The two antioxidants we tested, GTE and NDGA, exerted the largest effect on lifespan in manual survival assays (Figure 1). GTE extended lifespan in two species, *C. elegans* (two of three strains) and *C. tropicalis* (all three strains) (Figure 1a). The effect on lifespan was more pronounced in *C. elegans*, with mean lifespan extended >15% in both MY16 and JU775. In the laboratory *C. elegans* strain N2, we observed lifespan extension via GTE in only one lab, and the pooled results across labs was insignificant. We also observed lifespan extension with GTE in *C. tropicalis*, with all three strains tested showing small but significant increases in mean lifespan; the largest effect was in JU1630 (~11% increase in mean lifespan). In contrast, we found that all *C. briggsae* strains exhibited a significant decrease in survival when exposed to GTE, with mean lifespan decreasing by 10-19%. Overall, we observe a strong species-specific effect on longevity with GTE exposure.

We found that NDGA also impacted *Caenorhabditis* longevity in a species-specific manner (Figure 1b). In *C. elegans*, much like with GTE, we measured an 8-16% increase in the mean lifespan of strains MY16 and JU775, while N2 had a similar, albeit insignificant, lifespan increase. The effect of NDGA on lifespan was more variable than we recorded for GTE as we observed a significant decrease in survival in five of six *C. briggsae* and *C. tropicalis* strains.

### Antioxidants that decrease survival in certain genetic backgrounds extend lifespan in automated *C. elegans* Lifespan Machine assays

To increase throughput, we have implemented ALMs^31^ (Supplemental Figure 1, Phase Two) at all three CITP sites. The ALM is a lifespan analysis platform built on flatbed scanner technology that enables life-long imaging of animals at one-hour intervals, increasing both throughput and temporal resolution for data sampling. Our previous testing of compounds NP1, resveratrol, propyl gallate, thioflavin t, and α-ketoglutarate revealed that most ALM trials recapitulated outcomes from manual plate-based assays, although we did identify light sensitivity of some compounds as a factor that could change outcome ^5^. During preliminary experiments on ALMs we found that both GTE and NDGA increased lifespan in strains that also reported decreased lifespans in manual assays. More specifically, for GTE assays on the ALM, all *C. briggsae* strains exhibited increased survival compared to the control. For NDGA treatment, four of the six *C. briggsae* and *C. tropicalis* strains exhibited positive lifespan results on the ALM. Difference in outcome might be the consequence of increased light exposure on the ALMs^32^, as light is known to induce photooxidative stress^33^. ALM protocols also involve fewer potentially damaging animal transfers, and feature no exposure to freshly treated compound plates in mid-to late-life, in contrast to manual assays^34,35^. Our highly replicated data, however, emphasize that the methodological approach to lifespan determination is a factor in experimental outcome.

### GTE and NDGA impact on swimming ability is strain-specific

With the goal of maintaining health over the lifetime, compounds that slow the age-related decline in swimming ability are of particular interest to the CITP. We previously showed that swimming ability as a measure of locomotion with compound treatment in *Caenorhabditis* does not always correlate with lifespan, and thus swim locomotion is an assay with potential to identify compounds that may improve healthspan independent of lifespan (CITP, in preparation). Because both GTE and NDGA induced strong species-specific effects on lifespan, we addressed whether these compounds could improve locomotion.

We found that with GTE treatment, all *C. elegans* strains showed an improvement in swimming ability (Figure 2a). The improved locomotion effect was age-dependent in MY16 and JU775 (8-9% increase in mean swim score at old age as compared to the control, respectively), while N2 showed an overall robust improvement in locomotion (25% increase at young age, 146% increase at old age). The *C. tropicalis* strains (in which GTE improved lifespan), exhibited strain-specific responses in swimming ability: strain JU1630 showed no difference in locomotion with GTE treatment, while strain QG834 exhibited a small but significant decrease in swimming ability (2-4% decrease with age). JU1373 exhibited a strong age-related response in which GTE robustly decreased swimming ability at young age (14% decrease) but improved swimming ability during old age (16% increase). The effect of GTE on swimming ability in *C. briggsae* was minimal but strain-specific as we recorded decreases in swimming ability at young age in strains AF16 (23% decrease) and ED3092 (13% decrease), but a small increase at old age in HK104 (6% increase).

We also tested the effect of NDGA on locomotion in our panel of *Caenorhabditis* strains. With NDGA treatment, swimming ability improved in five of six *C. elegans* and *C. briggsae* strains, but none of the *C. tropicalis* strains (Figure 2b). The effect in *C. elegans* was observed in two strains, with N2 gaining a general increase (11-13% increase in mean swimming with age) and with JU775 exhibiting a reduction in the age-related decline of swimming ability (6% increase at old age). Interesting, all three *C. briggsae* strains showed an age-dependent improvement of locomotion at either young age (ED3092, 6% increase), or old age (HK104 and ED3092, 19-21% increase, respectively), irrespective of their reduced lifespan with NDGA treatment. *C. tropicalis*, which showed either no effect or a negative lifespan effect in response to NDGA, had no difference in swimming ability with NDGA treatment.

Overall, our tests underscore that: 1) longevity and locomotory health are not well correlated; 2) interventions can elicit marked differences in locomotory healthspan, even in the absence of longevity changes.

### GTE reduces oxidative stress resistance in some *Caenorhabditis* strains

We next assayed oxidative stress resistance with GTE treatment across our panel of strains because GTE exerted the most widespread effect on longevity and locomotion of the two antioxidants tested. In seven of the nine strains, GTE treatment reduced the animal’s ability to resist oxidative stress for at least one age tested (Figure 3). This effect was most prominent in the *C. briggsae* strains, which exhibited significant decreases in survival at all ages tested (ranging from a 17% decrease in mean survival in old AF16 to a 65% decrease in young ED3092). Previous exposure to a mild stressor can result in a more robust response to future stressors, a process known as hormesis^36^. Continuous exposure to an antioxidant may subsequently reduce reactive oxygen species that promote an organism’s ability to mount an enhanced oxidative stress response when removed from the compound. While GTE does have antioxidant properties^37^, the inability to resist oxidative stress may reflect an anti-hormetic effect. The only exceptions to the failure of GTE to increase oxidative stress resistance were in *C. tropicalis* JU1373, which exhibited an age-dependent increase in oxidative stress survival (22% increase at old age), and JU1630, with no effect at either age. *C. elegans* JU775 also had an increase in oxidative stress survival as compared to the control during old age (35% increase), though this increase was paired with a decrease in oxidative stress survival at young age (31% decrease).

We have found ALM-based thermotolerance assays to be challenging to reproduce across labs (CITP, in preparation). Still, as we continue to assess how broadly across compounds health measures are impacted, we tested thermotolerance consequent GTE treatment. As we previously observed, there was a high amount of variability within our dataset that made it difficult to identify significant effects on thermotolerance, but our results suggest that GTE may protect against thermosensitivity as most of the strains exhibited lifespan extension with GTE treatment (Supplemental Figure 4).

## Discussion

Aging is a complex trait with a broad range of observed outcomes. The inter-individual variability in outcome necessitates studies of significant size and replicate number to reliably identify anti-aging interventions. Because of the necessary study sizes, and the need to follow individuals over their lifetime, initial characterization of compound effects on aging is frequently performed in short-lived animal models. The oft-unspoken assumption is that the near universality of phenotypic aging in multi-cellular animals reflects shared underlying causes of aging, and insights gained in model systems will be generalizable across taxa. However, genetically identical individuals in the same environment can exhibit strikingly different aging trajectories. Whether that variability in outcome simply reflects differences in aging symptoms due to unidentified environmental differences or stochastic biological events or, importantly, if those differences reflect different causes of aging experienced by individuals is not known. The CITP attempts to account for issues of inter-individual differences and stochastic events by testing compounds at a large experimental scale using standardized protocols^34,35^, replicating experiments at three independent sites, and statistically partitioning the variance to determine the sources of variability (Supplemental Table 1 for this study). Additionally, the CITP tests compound interventions across a genetically diverse panel of *Caenorhabditis* strains and species to identify interventions that are effective independent of genetic background because they act through a widely conserved aging mechanism. The large number of animals studied, and the numbers of independent replicates in CITP assays, increase confidence that reported outcomes observed reflect true biology. Outcomes also underscore that few compounds may be expected to exert overarching impact on lifespan and mobility health. When such compounds are identified, a concerted effort to move them into to mammalian translation pipeline should be mounted.

### The CITP workflow

The CITP workflow needs to account for the multiplicative effects on labor needs due to replication of lifespan analyses at the three CITP sites, and the coverage across a genetic diversity panel. As such, CITP lifespan analyses are relatively labor-intensive, even when using short lived nematode models. The addition of health measures to the workflow necessitated further streamlining of CITP workflow (see Supplemental Figure 1). In the restructured CITP workflow we evaluate a compound’s ability to extend lifespan and/or healthspan across *Caenorhabditis* species and strains (Supplemental Figure 1). The pipeline involves preliminary testing in a single lab to narrow down the dosage range (phase one) and target the optimal dose across strains (phase two) before replicating lifespan results across labs (phase three) and evaluating the effect of robust positive compounds on healthspan (phase four). We found that compounds that did not look promising in the preliminary testing would not benefit from further investigation and have since introduced early exit points in the workflow whereby compounds without positive lifespan results are published and dropped from the pipeline ^6–8^.

### Examples of interventions that exert varied effects across species

As a group dedicated to identifying highly reproducible pharmacological interventions that extend lifespan, promote fundamental functionality such as locomotory ability, or both, with efficacy that applies over a genetically diverse *Caenorhabditis* test set, the CITP has first focused on testing compounds published to be effective in extending lifespan in nematodes or other model systems^4–7^. The five additional compounds we studied here, 17α-estradiol, acarbose, green tea extract, nordihydroguaiaretic acid, and rapamycin, are representative of those interest class interventions. We evaluated the five compounds for longevity modulation across a genetically diverse test set and pursued two of the most potent promoters of longevity, GTE and NDGA, for impact on locomotory health and stress resistance. We observed that the antioxidants GTE and NDGA modestly extend *Caenorhabditis* lifespan in a species-specific manner (Figure 1). Additionally, the antioxidants exhibited differences according to survival assay protocol, with observed decreased survival for certain genetic backgrounds in manual survival assays contrasting with extended lifespan as determined on the automated *C. elegans* Lifespan Machines (Figure 1 and Supplemental Figures 2–3). GTE and NDGA confer strain specific impact on swimming ability (Figure 2), but GTE reduces oxidative stress resistance in some *Caenorhabditis* strains (Figure 3), suggesting that GTE mechanistic targets and the underlying cause(s) of diminished health for these assessments may reflect different processes. While lifespan and health can certainly be uncoupled, and both are plausible targets for intervention, this study combined with previous observations underscores the complex challenge to finding universal lifespan and healthspan extending interventions.

### Implications for identifying compounds that slow aging using C. elegans

Our current study holds implications for future study design with chosen methodology potentially complicating identification of anti-aging compounds. For example, the differences observed in compound lifespan effects for manual and automated longevity studies for GTE and NDGA potentially reflect environmental differences between assay types that can profoundly alter compound longevity effect. The differences in measured effects may be easy to explain, for example GTE has been shown to be protective under photo-damage^38^, and the frequent illumination during automated lifespan analysis could induce photo-damage. Additionally, previous studies have concluded that the most abundant catechin in green tea, epigallocatechin gallate, didn’t extend lifespan under normal laboratory conditions, but did under stress conditions^23^, consistent with our observations if ALM assays are mildly stressful. Consistent with that interpretation, we previously found that under CITP protocols, measured lifespans for untreated animals are typically shorter in automated analyses relative to manual assays^5^. But the question remains; what environment is the most informative for translation to subjects outside of a controlled experimental environment? Additionally, the disconnect between lifespan and health measurements, and the genetic background dependency of those effects suggests additional complications that need to be addressed in a screening experimental design. In total, against a small but expanding test set, we observe that compound genetic background and assay type can give rise to differences in lifespan evaluation and health assessment^4,5^.

Stepping back to offer some perspective, it is likely unrealistic to expect many blockbuster aging interventions using broad based outcome evaluation. Nonetheless, the pursuit of the particular compounds that can positively move outcomes across genetic backgrounds and via multiple measures of heath might be the critical strategy that spotlights priority interventions for mammalian testing.

### Conclusion

Our study of five interventions (NDGA, GTE, 17α-estradiol, acarbose, and rapamycin) underscore the complexities of assessing biological outcomes of candidate aging interventions. Using our standardized protocols, we find that the antioxidants GTE and NDGA extend *Caenorhabditis* lifespan in a species-specific manner. GTE and NDGA tests also revealed some assay-specific outcomes—in certain genetic backgrounds we found decreased survival in manual longevity assays, whereas we measured extended lifespan when we determined outcomes using the automated *C. elegans* Lifespan Machines. GTE and NDGA affected swimming ability in a strain-specific manner, and GTE lowered oxidative stress resistance in some *Caenorhabditis* strains. Lifespan and healthspan appear uncoupled. Overall, our findings on this test set of interventions underscore how impactful genetic background, selected health assay, and protocol details are in the assessment of intervention effects. Interventions that meet the high bar of efficacy across a broad range of genetic backgrounds and across multiple experimental approaches may prove the exception, but such capacity would establish definitive priority for testing in mammalian models.

## Materials and methods

### Strains

All natural isolates used were obtained from the *Caenorhabditis* Genetics Center (CGC) at the University of Minnesota: *C. elegans* N2, MY16, and JU775; *C. briggsae* AF16, ED3092, and HK104; *C. tropicalis* JU1630, JU1383, and QG834. Worms were maintained at 20°C on 60 mm NGM plates seeded with *Escherichia coli* OP50-1.

### Interventions

Compound intervention treatments were performed as previously described^4^. The compounds used include green tea extract (LKT Laboratories, Inc. G6817, lot #2595901), nordihydroguaiaretic acid (Sigma-Aldrich 74540, lot #BCBQ4489V), α-estradiol (Sigma-Aldrich E8750, lot #016M4175V), rapamycin (LC laboratories R-5000, lot #ASW-135, and acarbose (Sigma-Aldrich A8980, lot #MKBS1059V0). Compounds were obtained as solids and dissolved in either water or DMSO (dimethyl sulfoxide) to obtain stock solutions, with either water or DMSO used to treat control agar plates. DMSO stock solutions (both compound and control) were then further diluted with water to create working solutions to allow for even distribution across the agar plate while maintaining a final concentration of 0.25% DMSO. Agar plates were treated with compound stock solutions such that the final volume was assumed equal to the volume of the agar.

### Manual lifespan assay

Per the previously published CITP standard operating procedure^4^, synchronized populations were generated via timed egg lays on 60 mm NGM plates. At day one of adulthood, 50 worms were transferred to 35 mm NGM plates containing 51 μM FUdR and compound intervention (or the solvent control). Worms were then transferred to fresh plates and scored as alive or dead on day two and five of adulthood for *C. elegans* and *C. briggsae*, or day two and four of adulthood for *C. tropicalis*. Thereafter, worms were transferred once weekly and scored every Monday, Wednesday, and Friday until dead. Death was defined as a lack of response when stimulated with a platinum wire.

### Automated lifespan assay

ALM assays were performed as previously described^5,32,34^, based on modification of the protocols published for the Lifespan Machine^31^. Briefly, worms were age synchronized and transferred to intervention plates as described above. One week post egg lay, animals were transferred to 50 mm tight-lidded, intervention treated, modified NGM plates containing 51 μM FUdR and 100 μM nystatin and loaded onto the ALMs. Scanner data was collected and analyzed using the Lifespan Machine software (https://github.com/nstroustrup/lifespan; ^31^ and strainspecific posture files^5^).

### Swimming ability assay

Swimming ability was measured per the standard CITP protocol (CITP, in prep;^39^), using the *C. elegans* Swim Test system (CeleST)^40,41^. Worms were age-synchronized and exposed to compound intervention during adulthood as described above, until swimming measurements were collected at ages 6 and 12 of adulthood (*C. elegans* and *C. tropicalis*), or ages 8 and 16 of adulthood (*C. briggsae*). Videos were processed using the CeleST software (https://github.com/DCS-LCSR/CeleST).

### Thermotolerance assay

The ability for animals to withstand heat stress was measured as previously published (CITP, in prep), utilizing a modification of the ALM protocol. Worms were synchronized and aged as adults on compound intervention plates, as stated above, until the desired testing age (adult day 6 and 12 for *C. elegans* and *C. tropicalis*, days 8 and 16 for *C. briggsae*). At this time, animals were placed onto 50 mm plates with modified NGM containing 100 μM nystatin, without FUdR or compound intervention, at a density of 70 worms per plate. Plates without lids were then transferred to Automated Lifespan Machines in an incubator set to 32°C and 50% humidity. Scanner data were collected using an increased scan speed and reduced resolution to provide proper temporal resolution, needed as a result of the shortened survival under these conditions. Images analyzed using the Lifespan Machine software (https://github.com/nstroustrup/lifespan)^31^ and strain-specific posture files ^5^ and deaths were validated by hand.

### Oxidative stress resistance assay

Resistance to oxidative stress was measured per the standard CITP procedure (CITP, in prep). Worms were prepared and aged as adults on interventions as mentioned above, with the same ages tested as in the swim test and thermotolerance assays. At the desired age, animals were transferred at a density of 70 worms per plate to 50 mm tight-lidded plates with modified NGM containing 40 mM paraquat (or methyl viologen dichloride, from Sigma-Aldrich), 51 μM FUdR, 100 μM nystatin. Plates were then transferred to ALMs at 20°C and scanner data collection, processing, and analyzing was done with the same methodology mentioned for the thermotolerance assay.

## Acknowledgements

We acknowledge the members of the Lithgow, Driscoll and Phillips labs for helpful discussions. We thank the CITP Advisory Committee and Ronald Kohanski (National Institute on Aging) for extensive discussion. We thank Asher Cutter, Marie-Anne Félix, and Christian Braendle for providing strains that they had directly collected. Additional strains were provided by the CGC, which is funded by NIH Office of Research Infrastructure Programs (P40 OD010440). This work was supported by funding from National Institutes of Health grants (U01 AG045844, U01 AG045864, U01 AG045829, U24 AG056052) and the Glenn Foundation for Medical Research and the Larry L. Hillblom Foundation.

## Tables

**Supplemental Table 1.** Comparison of reproducibility of manual lifespan assays within and between labs. Variance estimates were estimated using a general linear model, as previously described^4^. Bold entries represent categorical summation of the component numbers presented immediately beneath. Individual variation represents variability unassignable to a specific source of variance. Discrepancies between categorical totals and constituent sub totals represent deviations due to rounding.

| Source of Variation | Manual Assays |
| --- | --- |
| **Genetic Variation** | **22.5** |
| Among species | 6.8 |
| Among strains w/in species | 11.8 |
| Species x Compound | 3.6 |
| Strain x Compound | 0.3 |
| **Reproducibility Among Labs** | **3.6** |
| Among labs | 2.9 |
| Lab x species | 0 |
| Lab x strain | 0.7 |
| Lab x compound | 0.1 |
| **Reproducibility Within Labs** | **17.4** |
| Among trials | 0.5 |
| Among experimenters w/in trials | 2.9 |
| Among plates w/in experimenters | 14.0 |
| **Individual variation** | **56.5** |
| **Total** | **100** |
| Total number of observations | 73,003 |

## Supplemental Figures

**Supplemental Figure 1:**
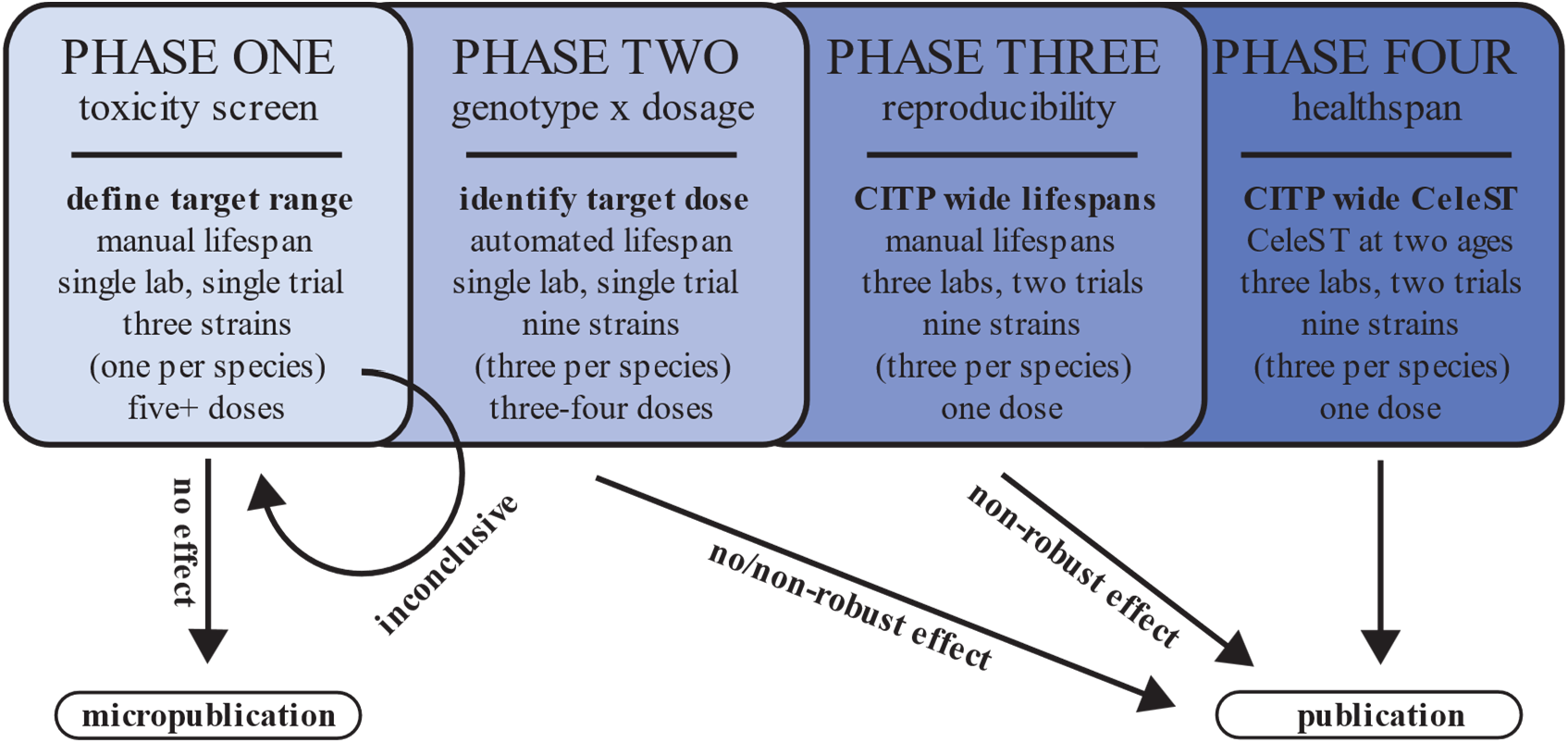
The CITP workflow for identifying compounds that robustly extend lifespan and/or healthspan across a genetically diverse set of *Caenorhabditis* species and strains. The pipeline involves screening for toxicity and positive hits (phase one), identifying a target dose (phase two), replicating lifespan effects across labs (phase three), and evaluating the effects of robust hits on healthspan (phase four). Exit points that result in publication after each phase were introduced to account for compounds that failed to exhibit benefits warranting movement to the next phase.

**Supplemental Figure 2:**
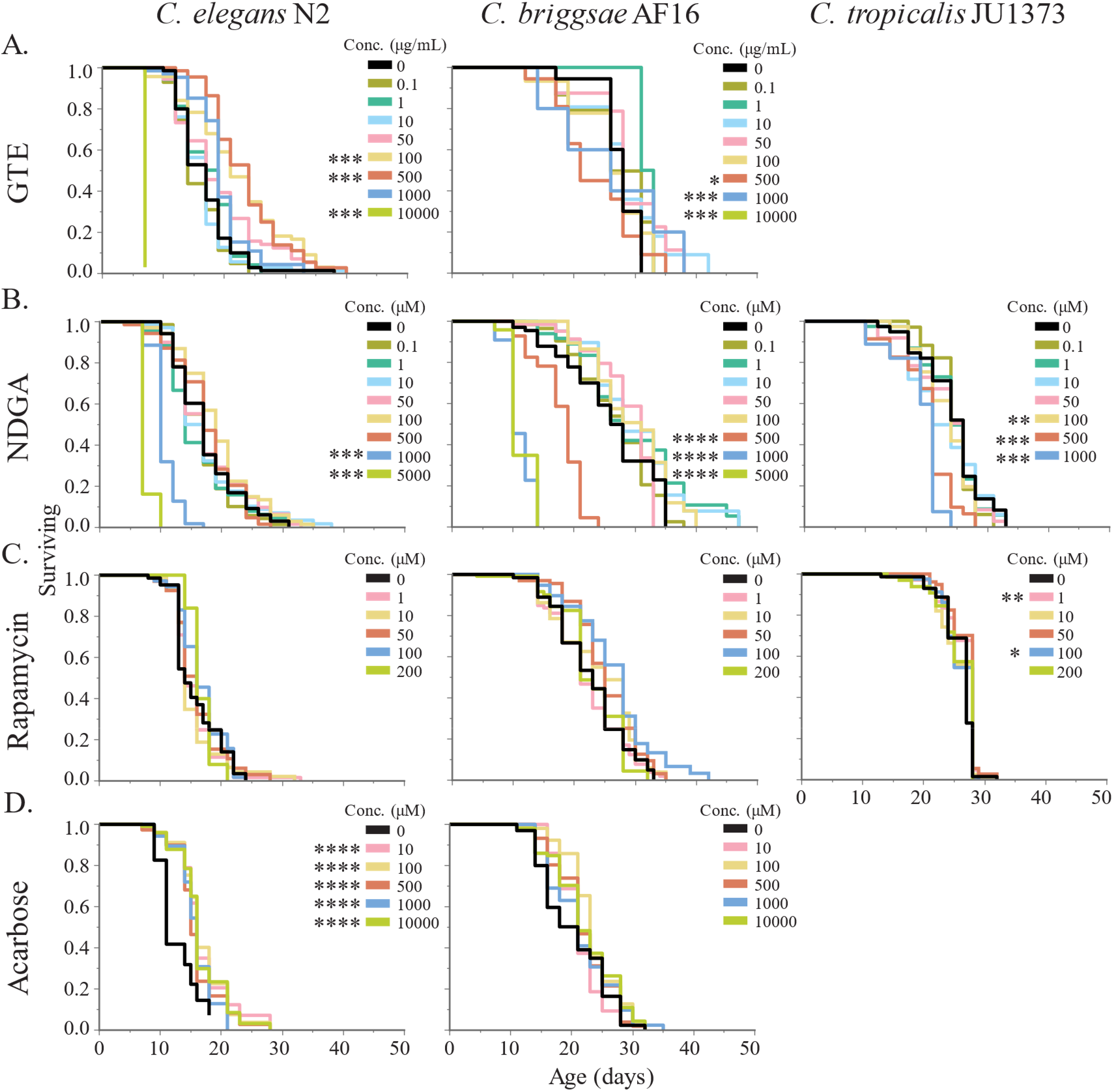
Data on initial dosing studies conducted prior to full CITP testing. Kaplan-Meier curves showing effect of adult exposure to multiple concentrations of (a) green tea extract, (b) NDGA, (c) rapamycin, (d), 17α-estradiol, and (e) acarbose on the lifespan of three strains across three *Caenorhabditis* species. One trial was conducted at one CITP testing site to find a target range of concentrations to be tested across nine strains. One strain was tested from each *Caenorhabditis* species: *C. elegans* N2, *C. briggsae* AF16, and *C. tropicalis* JU1373. The line represents the mean, and the bars represent the standard error of the mean. Asterisks represent *p*-values from the CPH model (except for AF16 analyses which used the general linear model because the CPH could not be calculated) such that \*\*\*\**p*<0.0001, \*\*\**p*<0.001, \*\**p*<0.01, and \**p*<0.05.

**Supplemental Figure 3:**
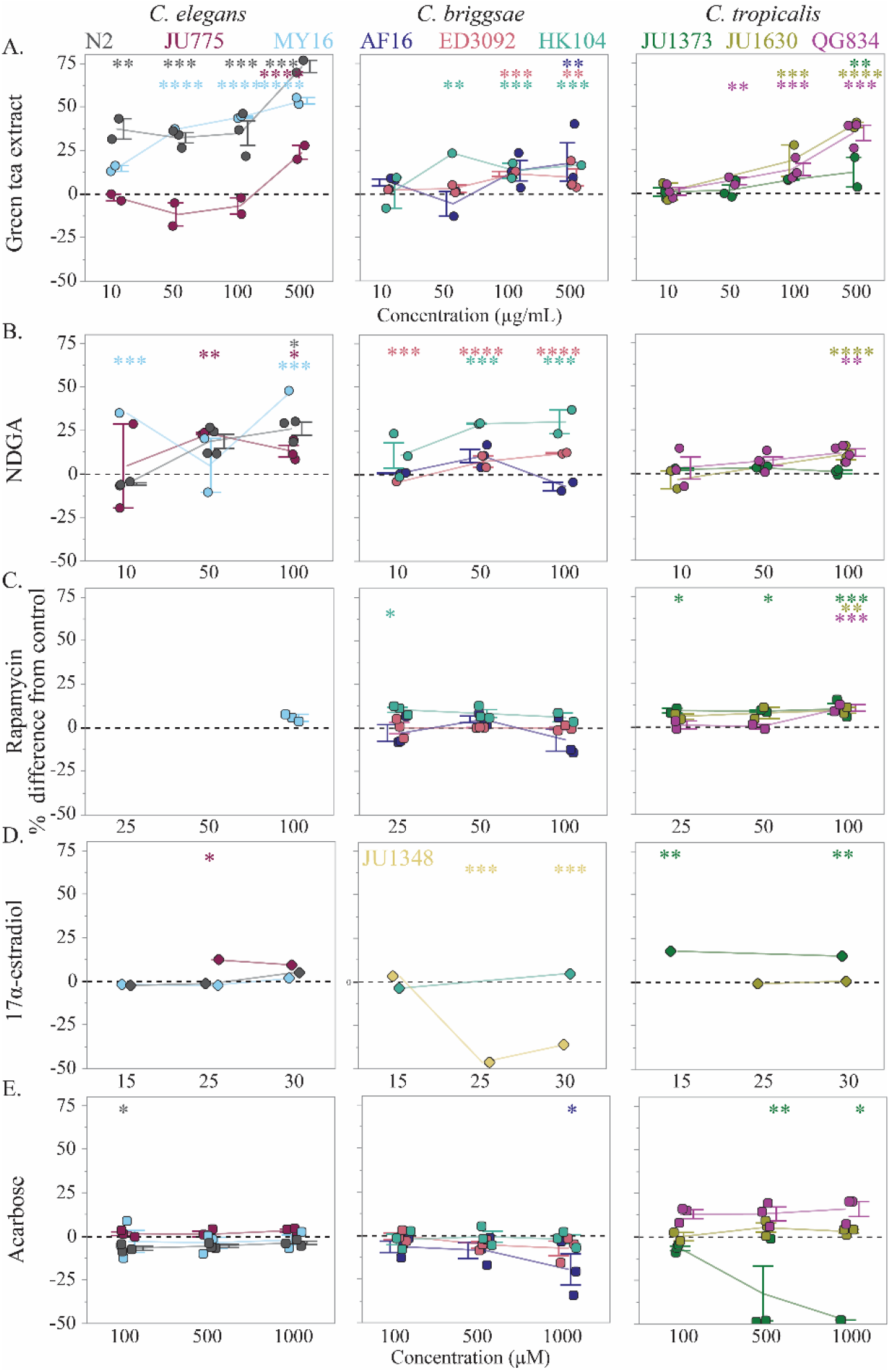
The effect of adult exposure to multiple concentrations of (a) green tea extract, (b) NDGA, (c) rapamycin, (d), 17α-estradiol, and (e) acarbose on the median lifespan of nine strains across three *Caenorhabditis* species. One trial was conducted using the Automated Lifespan Machines (ALMs) at one CITP testing site (square–Buck, circle–Oregon, diamond–Rutgers) in order to select a target dose for replicated, manual lifespan assays. Three strains were tested from each species: *C. elegans* N2 (gray), MY16 (light blue), and JU775 (burgundy), *C. briggsae* AF16 (navy blue), HK104 (teal), and ED3092 (pink) or JU1348 (pale yellow), *C. tropicalis* JU1373 (green), JU1630 (gold), and QG834 (purple). Each point represents the change in median lifespan from an individual trial plate (compound treated) relative to the specific control (vehicle only) conducted. The line represents the mean, and the bars represent the standard error of the mean. Asterisks represent *p*-values from the CPH model such that \*\*\*\**p*<0.0001, \*\*\**p*<0.001, \*\**p*<0.01, and \**p*<0.05.

**Supplemental Figure 4:**
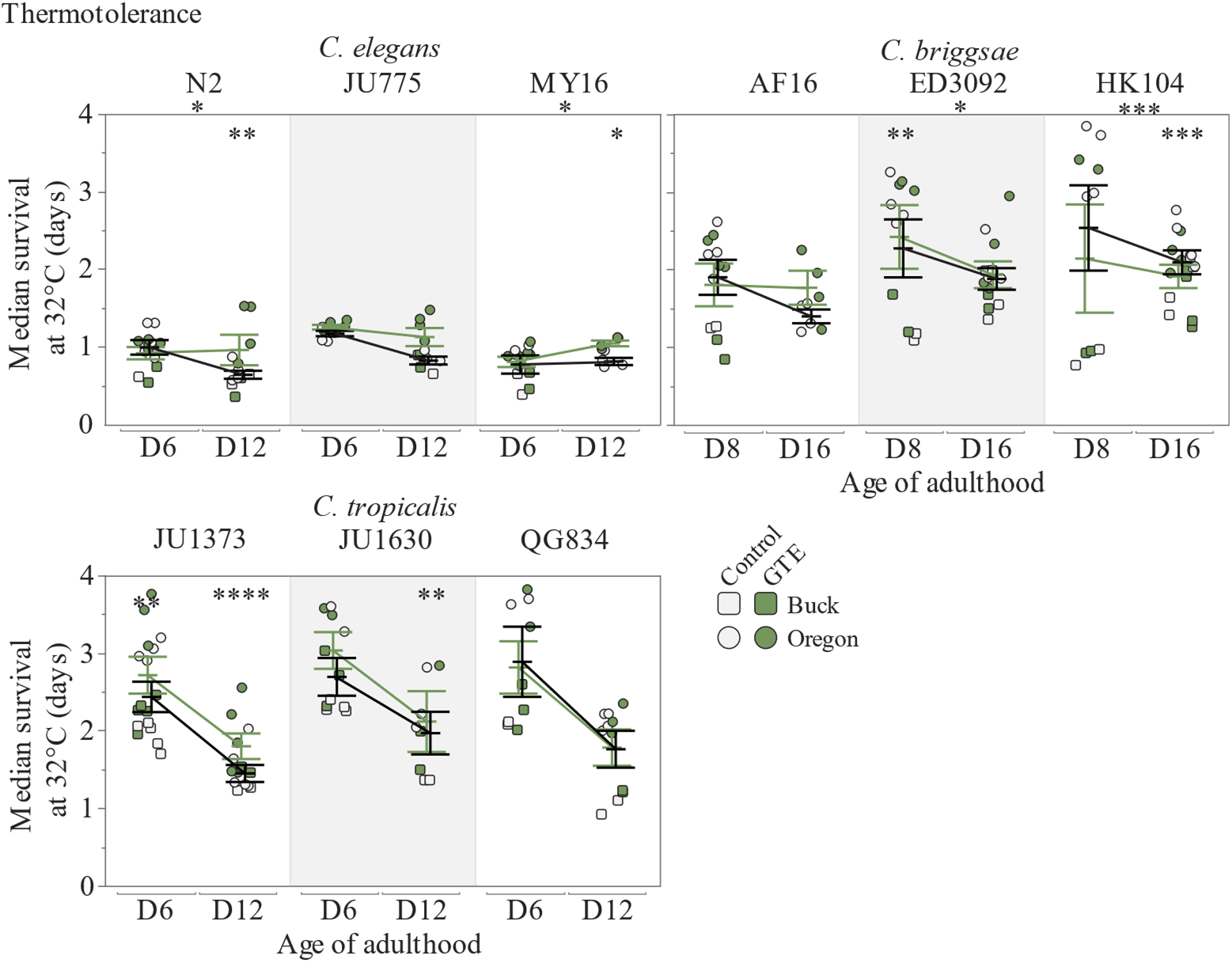
The survival of nine *Caenorhabditis* strains at 32°C after adult exposure to green tea extract. Thermotolerance was measured at day 6 and 12 of adulthood (*C. elegans* and *C. tropicalis*), or day 8 and 16 of adulthood (*C. briggsae*). Three strains were tested from each species: *C. elegans* N2, JU775 and MY16, *C. briggsae* AF16, ED3092 and HK104, and *C. tropicalis* JU1373, JU1630 and QG834. Dots represent the median survival of one plate replicate (gray–vehicle only control, green–500 μg/mL GTE). Replicates were completed at two CITP testing sites (circle-Oregon, square-Buck). Asterisks inside the plots represent *p*-values from the CPH model, and asterisks outside the plots represent *p*-values from the type III ANOVA indicating a significant compound by age interaction, such that \*\*\*\**p*<0.0001, \*\*\**p*<0.001, \*\**p*<0.01, and \**p*<0.05.

## Notes

### Competing Interest Statement

The authors have declared no competing interest.

